# The emergence, evolution, and diversification of the miR390-*TAS3-ARF* pathway in land plants

**DOI:** 10.1101/090647

**Authors:** Rui Xia, Jing Xu, Blake C. Meyers

## Abstract

In plants, miR390 directs the production of tasiRNAs from *TRANS-ACTING SIRNA 3* (*TAS3*) transcripts to regulate *AUXIN RESPONSIVE FACTOR* (*ARF*) genes, transcription factors critical for auxin signaling; these tasiRNAs are known as tasiARFs. This pathway is highly conserved, with the *TAS3* as the only one noncoding gene present almost ubiquitously in land plants. To understand the evolution of this miR390-*TAS3-ARF* pathway, we characterized homologs of these three genes from thousands of plant species, from bryophytes to angiosperms. Both miR390 and *TAS3* are present and functional in liverworts, confirming their ancestral role to regulate *ARFs* in land plants. We found the lower-stem region of *MIR390* genes, critical for accurate DCL1 (DICER-LIKE 1) processing, is conserved in sequence in seed plants. We propose a model for the transition of functional tasiRNA sequences in *TAS3* genes occurred at the emergence of vascular plants, in which the two miR390 target sites of *TAS3* genes showed distinct pairing patterns in different plant lineages. Based on the cleavability of miR390 target sites and the distance between target site and tasiARF we inferred a potential bidirectional processing mechanism exists for some *TAS3* genes. We also demonstrated a tight mutual selection between tasiARF and its target genes, and characterized unusual aspects and diversity of regulatory components of this pathway. Taken together, these data illuminate the evolutionary path of the *miR390-TAS3-ARF* pathway in land plants, and demonstrate the significant variation that occurs in the production of phasiRNAs in plants, even in the functionally important and archetypal miR390-*TAS3-ARF* regulatory circuit.

## Introduction

In plants, small RNAs (sRNAs) play crucial regulatory functions in growth and development, resistance to abiotic and biotic stresses, and reproduction (Chen 2009; Axtell 2013; Bartel 2009). Based on features such as their biogenesis and function, sRNAs are classified into two major groups, microRNAs (miRNAs) and small interfering RNAs (siRNAs). miRNAs are generated from precursor mRNAs that fold back to form double-stranded stem-loop structures, while siRNAs are produced from double-stranded RNAs (dsRNAs) biosynthesized secondarily by RNA-dependent RNA polymerase (RDR) (Axtell 2013). *Trans*-acting small interfering RNAs (tasiRNAs) are a special type of small RNAs found only in plants, so far. Precursor genes of tasiRNAs (*TAS* genes) are sliced in a miRNA-directed event, and the cleaved fragment is made double-stranded by RDR6; the resulting dsRNA is chopped by DICER-LIKE 4 (DCL4) into 21-nt siRNAs that map back to the precursors in a head-to-tail arrangement initiating from the miRNA cleavage site (Allen et al. 2005; Yoshikawa et al. 2005).

Among plant *TAS* genes, the most well-studied is *TAS3*; its transcript bears two target sites of miR390, generating tasiRNAs via the so-called “two-hit” mechanism (Axtell et al. 2006). The conserved, resulting tasiRNA is known as “tasiARF” as it targets auxin responsive factor (*ARF*) genes (Allen et al. 2005; Axtell et al. 2006). To date, there are two kinds of *TAS3* genes described in plants; one contains a single, centrally-located tasiARF, while the other generates two tasiARFs denoted as *TAS3*-short (*TAS3S*) and TAS3-long (*TAS3L*), respectively (Xia et al. 2015b). In *TAS3L*, only the 3′ miR390 target site is cleavable and this sets the phase of tasiRNA production, giving rise to the two in-phase tasiARFs (Allen et al. 2005; Axtell et al. 2006). The 5′ target site of *TAS3L* is usually non-cleavable because of the presence of a central mismatch (10^th^ position) in the pairing of miR390 and target site (Axtell et al. 2006). It serves as an important binding site of ARGONAUTE 7 (AGO7), a specialized protein partner of miR390 (Montgomery et al. 2008a). In contrast, both target sites of *TAS3S* are cleavable, and both can potentially initiate tasiRNA generation (Howell et al. 2007; Xia et al. 2012, 2015b). The single tasiARF of *TAS3S* is in near-perfect phasing to both miR390 sites as there is only 2-nt difference between the phase registers set by the two target sites (Xia et al. 2012, 2015b).

Auxin, a plant hormone, regulates seemingly every aspect of plant growth and development. The small class of ARF transcription factors can either activate or repress expression of downstream auxin-regulated genes through protein–protein interactions with auxin/indole-3-acetic acid (Aux/IAA) family members (Guilfoyle and Hagen 2007). Plant genomes contain ~10 to 30 *ARF* genes; for example, there are 23 members in the model plant Arabidopsis. ARFs are classified into three clades: ARF5/6/7/8 (Clade A), ARF1/2/3/4/9 (Clade B), and ARF10/16/17 (Clade C) (Finet et al. 2013). The *TAS3*-derived tasiARF specifically targets *ARF* genes of Clade B (ARF2/3/4). This miR390-*TAS3-ARF* pathway is of critical function in the regulation of plant growth and development, including leaf morphology, developmental timing and patterning, and lateral root growth (Garcia et al. 2006; Fahlgren et al. 2006; Adenot et al. 2006; Marin et al. 2010; Hunter et al. 2006). It was recently found that ARF3, with the transcription factor INDEHISCENT (IND), comprises an alternative auxin-sensing mechanism (Simonini et al. 2016). Loss-of-function mutants of AGO7, the specialized AGO partner of miR390, show varying degrees of growth and developmental disorders due to the malfunction of the tasiARF pathway (Yifhar et al. 2012; Zhou et al. 2013; Dotto et al. 2014). For example, maize *ago7* (*leafbladeless1*) plants have thread-like leaves lacking top/bottom polarity (Dotto et al. 2014), and Medicago *ago7* (*lobed leaflet1*) mutant plants displayed lobed leaf margins and extra lateral leaflets (Zhou et al. 2013).

All three components of the pathway, miR390, *TAS3*, and *ARFs*, are present in the oldest land plants, liverworts (Krasnikova et al. 2013; Finet et al. 2013). Interestingly, in bryophytes, the *TAS3* genes are different from those found in flowering plants. Although bryophyte *TAS3* genes also have two miR390 target, they generate tasiRNAs targeting not only *ARF* genes but also *AP2* genes (described, for example, in the moss *Physcomitrella patens*) (Axtell et al. 2007). Moreover, the bryophyte *ARF*-targeting tasiRNA is a different sequence compared to the tasiARF in flowering plants (Allen et al. 2005; Axtell et al. 2007). How and when this transition in *TAS3* gene composition occurred in the evolution of land plants is fascinating but unknown. We recently characterized ~20 *TAS3* genes in the gymnosperm Norway spruce (*Picea abies*), demonstrating diverse features of these genes distinct from those characterized in flowering plants (Xia et al. 2015a). In this study, we aimed to understand the evolutionary history of and critical changes in the miR390-*TAS3-ARF* pathway for the major lineages of land plants. We used > 150 plant genomes and the large dataset from the 1000 Plant Transcriptomes (1KP) project, in combination of additional sequencing data and computational approaches; these resources identified hundreds of *MIR390* genes and thousands of *TAS3* and *ARF* genes, across numerous plant species. From these data, we elucidated with high-resolution the dynamic nature of the evolutionary route of the miR390-*TAS3-ARF* pathway, revealing new regulatory features of the three critical components of the pathway.

## Results

### Gene identification from plant genomic and transcriptomic data

miR390-*TAS3-ARF* comprise a regulatory pathway highly conserved in plants. To maximize the possibility of characterizing the full diversity of the three main components of this pathway (miR390, *TAS3*, and *ARF* genes), we collected 159 sequenced plant genomes, ranging from liverworts to angiosperms, plus the 1KP data (Matasci et al. 2014). For the identification of *TAS3* genes, only genomic loci from sequenced genomes or transcripts (from 1KP data) containing at least one miR390 target site and one tasiRNA targeting *ARF* gene were considered valid for our analysis. Using bioinformatics tools and customized scripts (see Methods), we identified 374 *MIR390* genes from 163 plant species, 1923 *TAS3* genes from 792 species, and 2912 *ARF* genes (targets of tasiRNAs) from 934 species. We were unable to identify homologs of *MIR390* or *TAS3* genes in five algal genomes, consistent with the earlier conclusion that the miR390-*TAS3-ARF* pathway originated in land plants (Krasnikova et al. 2013).

To evaluate the evolutionary changes of three components of the pathway, we classified all the plant species into one of seven groups (liverworts, mosses, monilophytes or ferns, gymnosperms, basal angiosperms, monocots, eudicots); each group was considered independently in our subsequent assessments. Monocots and eudicots accounted for the two largest groups of species and yielded the vast majority of *MIR390* genes; many fewer were identified in the liverwort, monilophyte, and basal angiosperm groups (Fig. S1A). Similarly, most of the *TAS3* genes identified were from angiosperms, although there were many from gymnosperms as well (Fig. S1A).

We next examined variation in the length and GC content of *MIR390* and *TAS3*. Flanking sequences of 50 bp (5′ of the miR390 and 3′ of the miR390* for *MIR390*; 5′ of the 5′ miR390 target site and 3′ of the 3′ target site for *TAS3* genes) were included for these analyses. The length of the *MIR390* genes ranged from approximately 150 bp to 250 bp, with the *MIR390* copies in monocots significantly larger than those in gymnosperms (2.00E-05, t-test) and eudicots (4.00E-07, t-test) (Fig. S1B, at left). The GC content of *MIR390* genes was similar among different plant groups, with the exception of the eudicots in which they had a substantially lower GC content (1.40E-09, t-test) (Fig. S1C, at left). For the *TAS3* genes, their length was noticeably shorter in monilophytes, while significantly longer in gymnosperms than in monocots and eudicots (2.00E-16, t-test); in gymnosperms, there is an apparently bimodal distribution of lengths, possibly reflecting that there are two major types of *TAS3* genes of different length (Fig. S1B, at right). The GC content of the *TAS3* genes of two groups, the mosses and eudicots, was exceptional: the moss *TAS3* genes were of higher GC content (2.00E-16, t-test), while the eudicot *TAS3* genes had a much lower GC content (2.00E-16, t-test, Fig. S1C, at right).

In Arabidopsis, the proper execution of the miR390-*TAS3-ARF* pathway requires that miR390 is loaded into a specific and highly selective AGO partner protein, AGO7 (Montgomery et al. 2008a). AGO7 is an indispensable component of the pathway, and thus we also investigated the evolutionary history of AGO7. To complement our recent survey of AGO proteins that mainly focused on flowering plants (Zhang et al. 2015), our analyses here focused on AGO proteins from non-flowering plants. We identified 237 AGO protein sequences with ≥ 800 amino acids, and these were used for the construction of a phylogenetic tree in combination with AGO proteins from three representative angiosperms: *Amborella trichopoda, Oryza sativa*, and *Arabidopsis thaliana.* As previously documented (Vaucheret 2008; Mallory and Vaucheret 2010; Zhang et al. 2015), AGO proteins clustered into three major clades, AGO1/5/10, AGO2/3/7, and AGO4/6/8/9 (Fig. S2A). The AGO2/3/7 clade consisted of members all from vascular plants, except two moss AGO proteins (4_Pp3c17_350V3.1 from *Physcomitrella patens* and 4_Sphflax0148s0007.1 from *Sphagnum fallax*) (Fig. S2B); we interpreted this as an indication that the ancestor of the AGO2/3/7 clade likely separated from the AGO1/5/10 clade in mosses. Also, AGO7 was apparently not specified until the emergence of gymnosperms, as only gymnosperm and *Amborella* AGOs joined the eudicot AGO7 copies to form a subclade (Fig. S2B). These results suggest that the specific partner AGO of miR390, AGO7, emerged much later than the miRNA and the pathway, possibly to enable unique functions of the miR390-*TAS3-ARF* pathway in seed and flowering plants.

### The lower stem region of MIR390 is under strong selection for conservation

miR390 is one of the most ancient miRNAs, well conserved in land plants. During the course of evolution, *MIRNA* genes (i.e. the precursor mRNAs) are relatively labile, typically displaying conservation only in the sequences of the miRNA and miRNA* in the foldback region (Jones-Rhoades et al. 2006; Fahlgren et al. 2010; Ma et al. 2010). Indeed, in our analysis, the sequences of miR390 and miR390* were extremely conserved in land plants, as shown in the sequence alignment in Fig. 1A. Interestingly, in addition to the miR390/miR390* region, we identified another two regions of relatively high conservation in the precursors (Fig. 1A); these are the sequences forming the lower stem of the *MIR390* stem-loop structure (Fig. 1B). They displayed a substantially greater consensus, especially for the seed plants, than any other regions of the precursors except the miR390/miR390* duplex (Fig. 1A).

**Figure 1.**
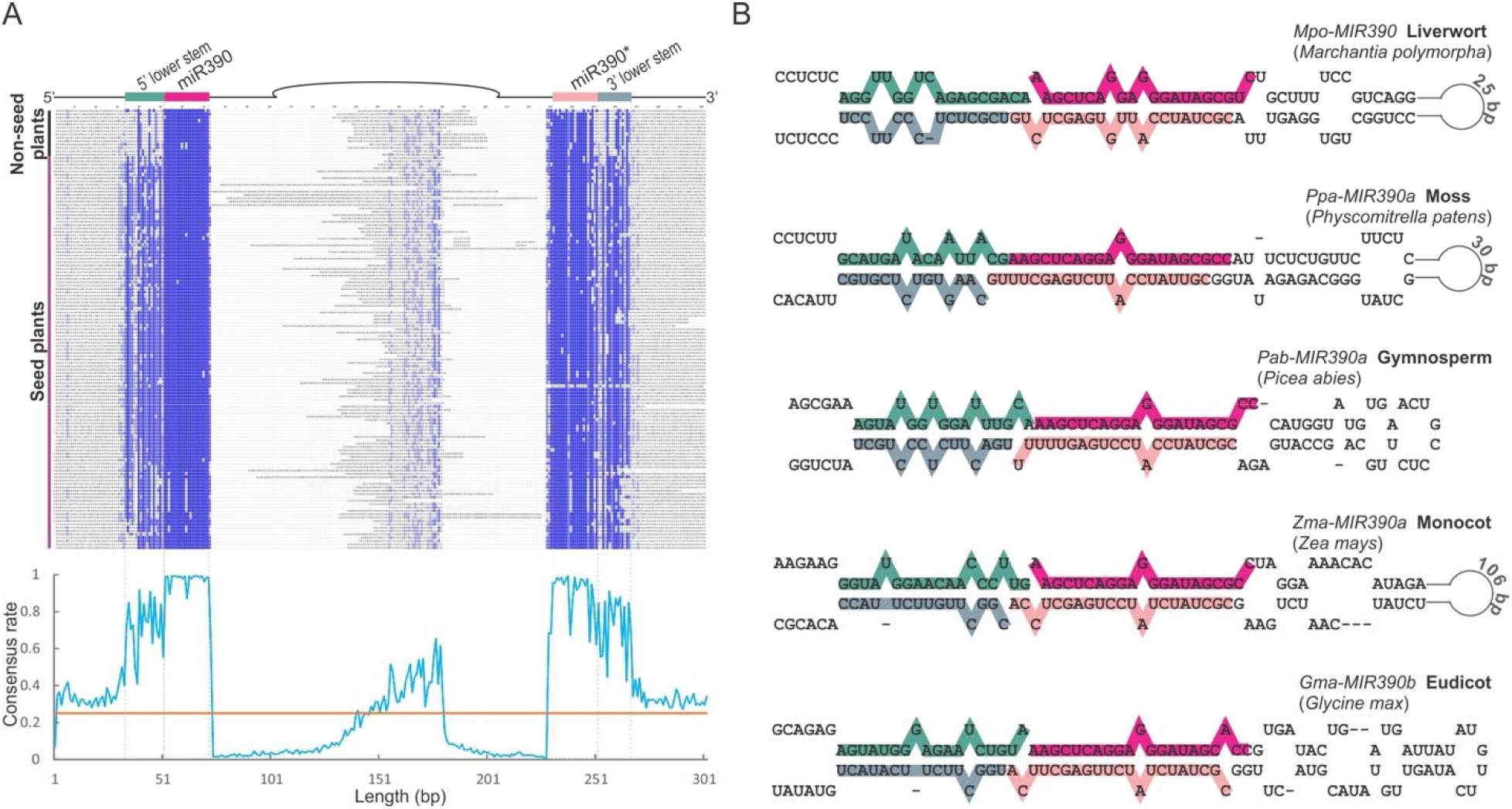
The lower stem region of *MIR390* is conserved in land plants. (A) Nucleotide sequence alignment of *MIR390* precursor genes (±50 bp before/after the miR390/miR390* region) with different sequence regions denoted above. The consensus rate of diversity of each position in the alignment is shown in the plot below with the orange line indicating the 25% level, since in a sequence randomized by neutral evolution, each nucleotide (A/U/C/G) would comprise 25% of each position. (B) Examples of stem-loop structures of *MIR390* precursor transcripts. The miRNA and lower-stem regions are indicated according to the colors shown in the top of panel A.

In plants, the release of a miRNA/miRNA* duplex relies on two sequential cuts by DCL1 in the *MIRNA* stem-loop precursors. These two sequential cuts are directional, either base-to-loop or loop-to-base. For base-to-loop processing, the first cut is defined by the distance from the miRNA/miRNA* duplex to a large loop at the base; the distance is usually ~15 nt (Werner et al. 2010; Song et al. 2010; Mateos et al. 2010). miR390 is one such base-to-loop-processed miRNA (Bologna et al. 2013). The conservation in the *MIR390* lower stem, exemplified in Fig. 1B, is likely to maintain the consistent distance of ~15 nt to ensure the accuracy of the first cut by DCL1 of the *MIR390* stem-loop precursor.

### TAS3 originated to regulate ARF genes

*TAS3* genes in bryophytes were firstly characterized in the moss *Physcomitrella patens*, consisting in that genome of a small family of six genes (Axtell et al. 2007; Arif et al. 2012). Many *TAS3* genes were subsequently described in mosses (Krasnikova et al. 2013). All known moss *TAS3* genes have similar sequence components: two miR390 target sites, a tasiRNA targeting *AP2* genes (tasiAP2), and a tasiRNA targeting *ARF* genes (tasiARF)(Axtell et al. 2007; Krasnikova et al. 2013). A *TAS3* gene was also identified in a liverwort *Marchantia polymorpha*, representing the most ancient extant lineage of land plants (Krasnikova et al. 2013). However, this *TAS3* gene was described to produce only a single tasiRNA of sequence similar to the moss tasiAP2. We found five *TAS3* genes from liverwort species in addition to that of *Marchantia polymorpha.* Sequence alignment of these six liverwort genes revealed the presence of another conserved region, aside from the two miR390 target sites and the previously-described tasiAP2, that could also produce an siRNA (Fig. 2A). Analyses of public sRNA data from *Marchantia polymorpha* showed that a highly abundant tasiRNA was produced from the anti-sense strand of this siRNA site. This tasiRNA (hereafter, “tasiARF-a1”) was predicted to target an *ARF* gene in *M. polymorpha*, with the cleavage of the target site confirmed by PARE analysis (Fig. 2A). While the previously-described tasiAP2 site is highly conserved, we were unable to validate its target interaction in *M. polymorpha* in which we attempted by combining whole genome target analysis with sRNA and PARE data. This is likely for several reasons: first, the corresponding tasiRNA was produced of low abundance; second, no *AP2* homolog was predicted as a target of the tasiRNA even using relaxed prediction criteria (alignment score ≤ 7); third, after checking *M. polymorpha* homologs of moss *AP2* genes that are validated targets of moss tasiARF-a1, we found no tasiAP2 target sites (data not shown). Therefore, the miR390-*TAS3* machinery likely originated to regulate *ARF* genes, and *not AP2* genes, unlike previous reports.

**Figure 2.**
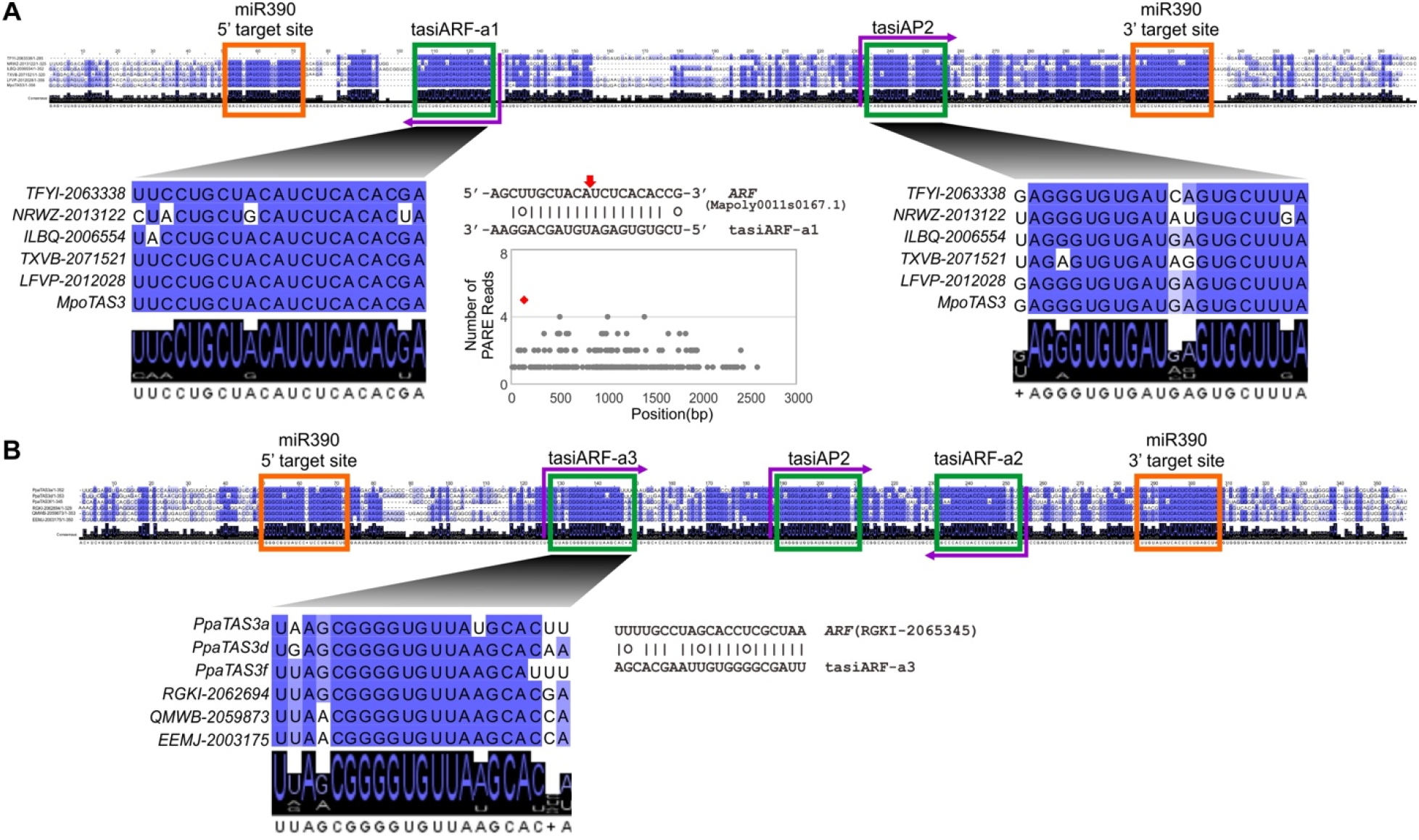
*TAS3* originated to regulate *ARF* genes in land plants. (A) Conserved motifs in *TAS3* transcripts from liverworts. Purple arrows indicate the encoded strand of tasiRNAs; the left-pointing arrow indicates that the functional tasiRNA is located on the anti-sense strand, and the right-pointing arrow indicates that the tasiRNA is on the sense strand. tasiARF-a1 encoded in liverwort *TAS3* genes targets *ARF* genes with the tasiRNA:target pairing (cleavage site marked with a red arrow) and validating, experimentally-derived PARE data shown in the middle. The red dot in the plot of PARE data marks the cleavage site directed by tasiARF-a1. (B) Conserved motifs in a few representative *TAS3* genes in mosses. Besides tasiARF-a2, previously reported to target *ARF* genes, another tasiRNA, denoted as tasiARF-a3, was predicted to target *ARF* genes. The tasiARF-a2 and tasiARF-a3 sequences are encoded in the anti-sense and sense strands of *TAS3* transcripts, respectively.

For the bryophytes, we identified a large number of *TAS3* genes including 67 genes from 36 moss species), in addition to the six liverwort *TAS3* copies described above. For the 67 genes, we built a multiple sequence alignment; from this, conserved sequence motifs, including the two miR390 target sites, and both tasiAP2 and tasiARF, were detected as previously reported and as observed in the mosses (Fig. S3A). In addition, we identified another tasiRNA site which is conserved only in a subset of the moss *TAS3* genes. Target predictions indicated that this tasiRNA may target *ARF* genes as well, and it is conserved in only a few members of the *TAS3* family, for instance, three *TAS3* genes of *P. patens* (*a/d/f*) encode this tasiRNA sequence, but *TAS3b/c/e* lack it (Fig. 2B and Fig. S3A). In contrast to the previously-identified tasiARF (tasiARF-a2, on the 3′ end) which was produced in the antisense strand and in phase with the 3′ miR390 target site, the newly identified tasiARF-a3 is located in the sense strand and in phase with the 5′ miR390 target site (Fig. 2B). These three tasiARFs in liverworts (tasiARF-a1) and mosses (tasiARF-a2, -a3) have no sequence similarity, originate from either strand of *TAS3* genes, and target different regions of *ARF* genes, consistent with independent origins.

The distribution of tasiARF-a2 and -a3 in moss *TAS3* genes is consistent with distinct evolutionary paths for these genes. To infer the possible evolutionary paths of *TAS3* in bryophytes, we constructed a phylogenetic tree using their *TAS3* genes. The phylogenetic tree (Fig. S3B) yielded three major classes: class I contained all six liverwort *TAS3* genes (tasiAP2 and tasiARF-a1); class II included moss *TAS3* genes containing tasiAP2 and tasiARF-a2; and class III comprised moss *TAS3s* with tasiAP2, tasiARF-a2, and tasiARF-a3. Intriguingly, class II is closer to liverwort *TAS3* genes (class I), indicating that class III *TAS3s* likely evolved after the appearance of the class II *TAS3* genes, which raises an interesting question of the origin of tasiARF-a3.

### The evolutionary path of TAS3 genes in land plants

*TAS3* genes found in seed plants are different from the bryophyte *TAS3s*. As summarized in Fig. 3A, two types of *TAS3* genes, *TAS3L* with two tandem tasiARFs and *TAS3S* with one tasiARF, have been previously characterized in gymnosperms and many angiosperms. Despite a similar arrangement of two miR390 target sites, the near-identical tasiARFs in *TAS3L* and *TAS3S* are distinct from moss tasiARFs (tasiARF-a1/a2/a3) in bryophyte *TAS3* genes, in terms of sequence, position and strand (Fig. 3A). These differences suggest a significant change occurred during *TAS3* evolution in land plants. To better understand when this change happened, we cataloged *TAS3* genes with tasiARFs; we found two *TAS3* genes from a lycophyte *Phylloglossum drummondii*, one with two tasiARFs (*Pdr-TAS3L*) and the other with a single tasiARF (*Pdr-TAS3S*) (Fig. 3B). The cDNA sequence of *Pdr-TAS3S* was too short to include the 5′ miR390 target site. We generated sRNA sequencing data which confirmed the phased generation of tasiARFs from *Pdr-TAS3L* (Fig. 3C). Both tasiARFs were predicted to target two *ARF* genes, found among the cDNA sequences from the same species (Fig. 3C). Therefore, we believe that this transformation of *TAS3* genes, and particularly the tasiARF transition, occurred after mosses and before or in lycophytes, perhaps with the emergence of vascular plants. Thus, we named all the *TAS3* genes producing these characteristic tasiARFs (i.e. not tasiARF-a1/a2/a3) as “vascular *TAS3*” genes.

**Figure 3.**
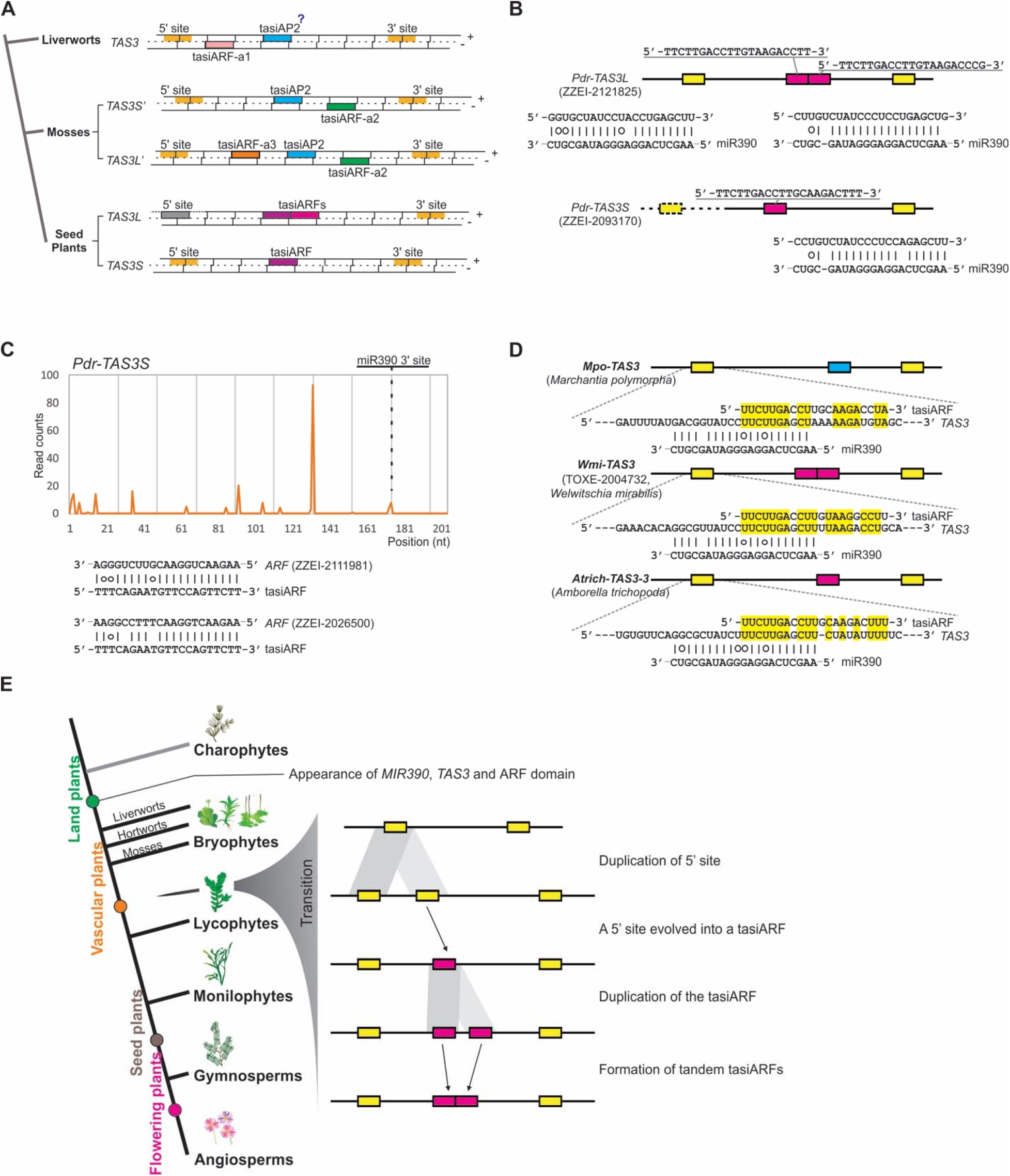
The inferred evolutionary progression of *TAS3* genes in land plants. (A) A summary of *TAS3* gene structures observed in land plants. Colored bars denote different features, as indicated; the grey 5′ miR390 site is not cleaved. The question mark “?” denotes that the function of tasiAP2 (targeting AP2 genes) could not be validated in liverworts. (B) Two *TAS3* gene structures found in the lycophyte species, *Phylloglossum drummodii.* (C) *TAS3* transcripts produce tasiARFs to regulate *ARF* genes in *Phylloglossum drummodii*. (D) tasiARF shows sequence similarity to the region partially covering the 5′ miR390 target site of cognate *TAS3* genes. Three representative *TAS3* genes from different species are displayed here. Identical nucleotides between tasiARF and the region partially covering the 5′ miR390 target site are highlighted in yellow. (E) An evolutionary model for the divergence of *TAS3* genes in land plants. The tasiARF sequence originated from the duplication of the 5′ miR390 target site and the *TAS3S* genes (with a single tasiARF) might be the ancestor of the *TAS3L* genes (with two tandem tasiARFs).

We next asked how the transition of *TAS3* genes happened. In other words, how was this signature tasiARF sequence generated in vascular plants. We compared the tasiARF sequence to available cDNA or genome sequences. We found the tasiARF sequence shared substantial sequence similarity to a region partially overlapped with the 5′ miR390 target site from the cognate *TAS3* gene in some species, as exemplified in a few *TAS3* genes shown in Fig. 3D. In a *TAS3* gene of the liverwort *Marchantia polymorph* (*Mpo-TAS3*), the tasiARF sequence has 15 nucleotides of identity with the 5′ miR390 target sequence, with an overlap of 11 nt. This sequence similarity is even greater in *TAS3* genes in a monotypic gymnosperm, *Welwitschia mirabilis* (*Wmi-TAS3*), and the basal angiosperm *Amborella trichopoda* (*Atrich-TAS3*) (Fig. 3D). This finding of sequence similarity is consistent with a hypothesis that the tasiARF was derived from the 5′ miR390 target site from *TAS3*.

We previously reported that the genome of the gymnosperm Norway spruce includes a large number of *TAS3* genes, of which many have non-canonical sequence features (Xia et al. 2015a). We extended this observation to other plant species, finding *TAS3* genes with varied motif structures in our large dataset (Fig. S4). For example, some have two 5′ or 3′ target sites due to short sequence duplications; some have two or three non-adjacent tasiARFs. We propose a model, consistent with these extant *TAS3* arrangements, for the tasiARF transition from bryophyte *TAS3* genes to vascular *TAS3* genes (Fig. 3E). In the first step, the 5′ miR390 target site of a bryophyte *TAS3* gene was duplicated through segmental duplication, as evidenced in a couple of gymnosperm *TAS3* genes. Next, the miR390 target site in the middle evolved into a tasiARF and was retained because of its essential function, yielding the short *TAS3* gene (*TAS3S*); after this, two tasiARFs in a single *TAS3* gene resulted from the duplication of tasiARF. Finally, the gap between the two tasiARFs was lost, forming a tandem repeat of tasiARFs, yielding the long *TAS3* gene (*TAS3L*) present in vascular plants. This series of steps is consistent with the *TAS3* variants present in plant genomes (Fig. S4).

### Distinct pairing patterns of two miR390 target sites

*TAS3* genes usually comprise a small gene family in plants. For instance, in bryophytes, only one *TAS3* gene was identified in *M. polymorpha*, and six *TAS3* copies in *P. patens*. For comparison, there are three *TAS3* copies in Arabidopsis, five in rice, and nine in maize – all vascular plants. Comparing across the 157 vascular plants with full-genome sequences that we utilized, we found that this size of the *TAS3* gene family is maintained across angiosperms, with most having fewer than ten *TAS3* genes and a mean of four genes (Fig. S5). This is in a sharp contrast to gymnosperms in which the *TAS3* family is substantially larger. The five gymnosperm species surveyed have at least 28 copies of *TAS3* genes, with the *Pinus taeda* encoding as many as 71 *TAS3* copies. Another noticeable feature of the vascular *TAS3* genes is that almost all of the species have both variants of *TAS3* genes (*TAS3L* and *TAS3S*) (Fig. S5), from which we infer that these two types of *TAS3* genes likely have non-redundant functions.

We next evaluated how essential sequence motifs of *TAS3* genes changed in vascular plants. We identified 3684 target sites of miR390 in 1847 vascular *TAS3* copies, including 1793 5′ sites and 1891 3′ sites. These 5′ and 3′ miR390 sites showed different patterns of pairing with miR390, of which sequence is highly conserved (Fig. 4A). In general, the majority of the 5′ sites encode a central, 10^th^-position mismatch, while the last four nucleotides of the pairing (18^th^ to 21^st^, relative to the 5′ end of miR390) are always mismatched in the 3′ target site (Fig. 4A). More specifically, the middle region (8^th^ to 12^th^ nucleotides) of the 5′ target site are of greater nucleotide diversity, with the 10^th^ position generally unpaired and the 11^th^ position is predominantly a G:U pair. In contrast, the 5′ five nucleotides (17^th^ to 21^st^, relative to the 5′ end of miR390) of the 3′ target sites vary substantially in sequence, with the last four (18^th^ to 21^st^) always unpaired with miR390. Noticeable is that the final nucleotide of the 3′ site (1^st^ relative to miR390) is not well conserved at all, maintained as a mismatch with miR390, unlike the 5′ site (Fig. 4A).

**Figure 4.**
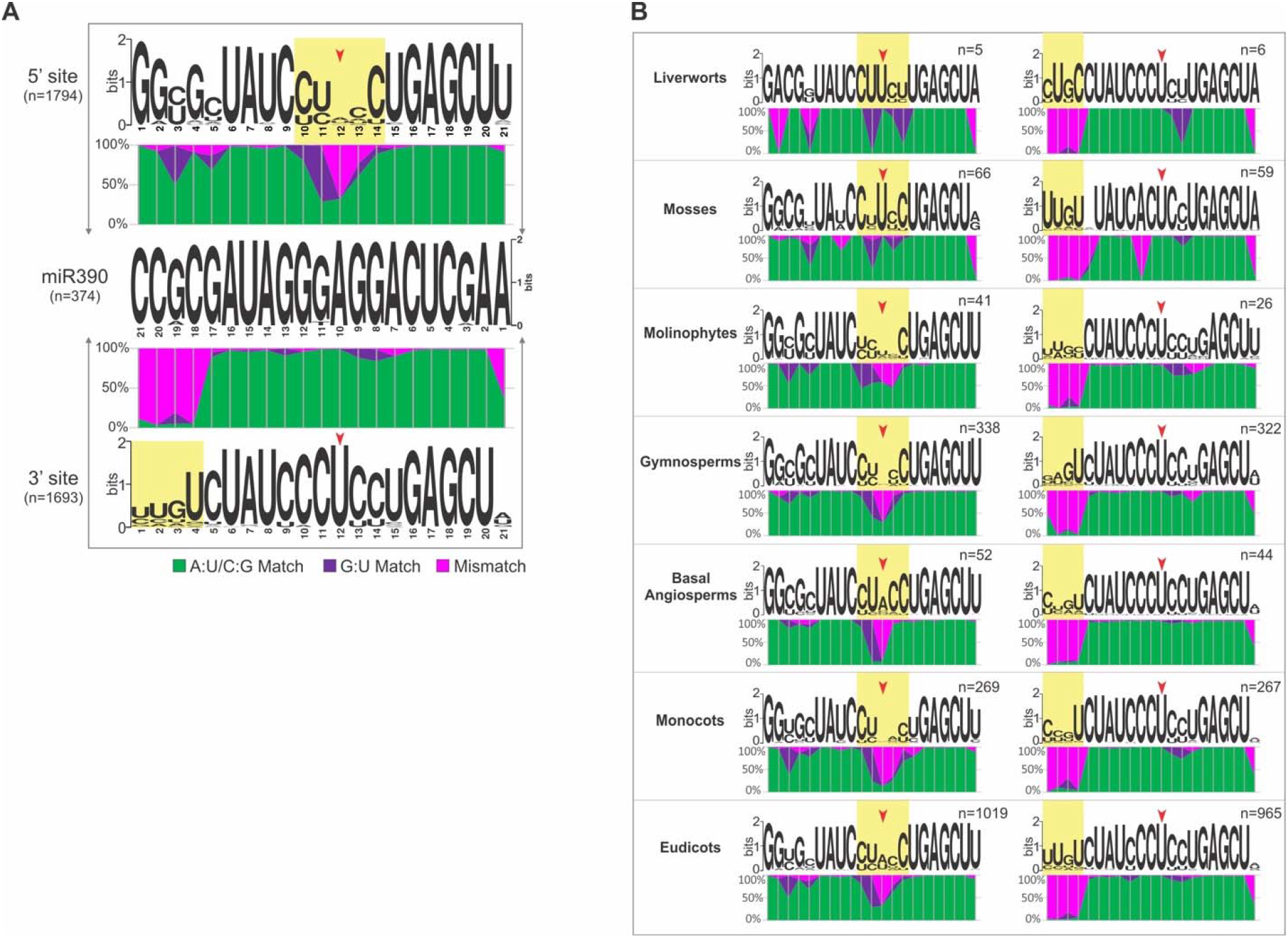
Pairing features and evolutionary variation of the two target sites of miR390 in *TAS3* genes. (A) Distinct pairing patterns of the two miR390 target sites in *TAS3* genes. Sequence logos were generated using WebLogo. Different nucleotide pairings at each position in the target site (compared to the highly conserved miR390 sequence in the middle) are indicated by different colors, with A:U/C:G matches denoted in green, G:U matches in purple, and all mismatches in pink. The red arrow marks the 10^th^ position, relative to the 5′ end of miR390. The yellow shading indicates regions of substantially imperfect pairing. The upper graph shows the 5′ target site of *TAS3*, the lower graph shows the 3′ target site; the number of sequences analyzed is indicated for each panel. (B) Variation in the pairing of the two miR390 target sites in *TAS3* genes in different species or lineages of land plants. The images are as described for panel A, but the left graph shows the analysis of the 5′ target sites of *TAS3*, and the right graph shows the analysis of the 3′ target sites.

To assess the history of diversification of the pairing between miR390 and its target sites in *TAS3* genes, we grouped all identified miR390 target sites according to the seven species lineages of land plants described above, and we generated similar plots to represent miR390-*TAS3* pairing. We observed substantial variation in pairing in the 5′ site, especially for the middle region (8^th^ to 12^th^ positions) (Fig. 4B). Interestingly, the position most important for AGO-mediated slicing, the 10^th^ position (of miR390) was always matched in bryophytes, yet in later-diverged species, the mismatch at this position appeared and seemed preferentially retained, as the proportion of mismatches gradually increased over plant evolution. This was particularly noticeable in the basal angiosperms and monocots in which there were almost no matched interactions at this position. For the 11^th^ position of the 5′ site, G:U pairing predominated in all the lineages in spite of a substantial portion of perfect G:C pairing at the 10^th^ position observed in Monilophytes (Fig. 4B). Regarding the 3′ site, its main features do not vary much among the groups, including the 5′ end mismatch region, the perfect match in the middle, and the high proportion of mismatches for the final nucleotide (except for the Monilophytes) (Fig. 4B).

### Evolutionary dynamic distances between tasiARFs and miR390 target sites

The tasiARF is another functionally essential component of the pathway of our investigation. To correctly generate the tasiARF, this siRNA needs to be in phase with a miR390 target site; in other words, the distance from the cleavage site of miR390 target site to the end of the tasiARF must be a multiple of 21 nucleotides. Therefore, we calculated the distances and evaluated their evolutionary changes from both 5′ and 3′ miR390 target sites to the tasiARF ends. Given that the tasiARF in vascular plants is distinct from tasiARF-a1/a2/a3 found in bryophytes, which themselves vary substantially, and given the large number of *TAS3* genes identified for vascular plants, we performed distance analyses only for vascular *TAS3* genes.

Overall, there was substantial variation in the tasiARF distances (5′-site → tasiARF and tasiARF → 3′-site) in all lineages of vascular plants, with the exception of the eudicots, in which the tasiARF → 3′-site distance of *TAS3L* and the 5′-site → tasiARF distance of *TAS3S* were highly consistent in length (Fig. 5A and B). For *TAS3L*, both distances were significantly shorter in the Monilophytes, but the gymnosperms had a much longer 5′-site → tasiARF region compared with other lineages (Fig. 5A). Monilophyte *TAS3S* also had a shorter 5′-site → tasiARF region, but the tasiARF → 3′-site distance was more or less similar to those of other lineages (Fig. 5B).

**Figure 5.**
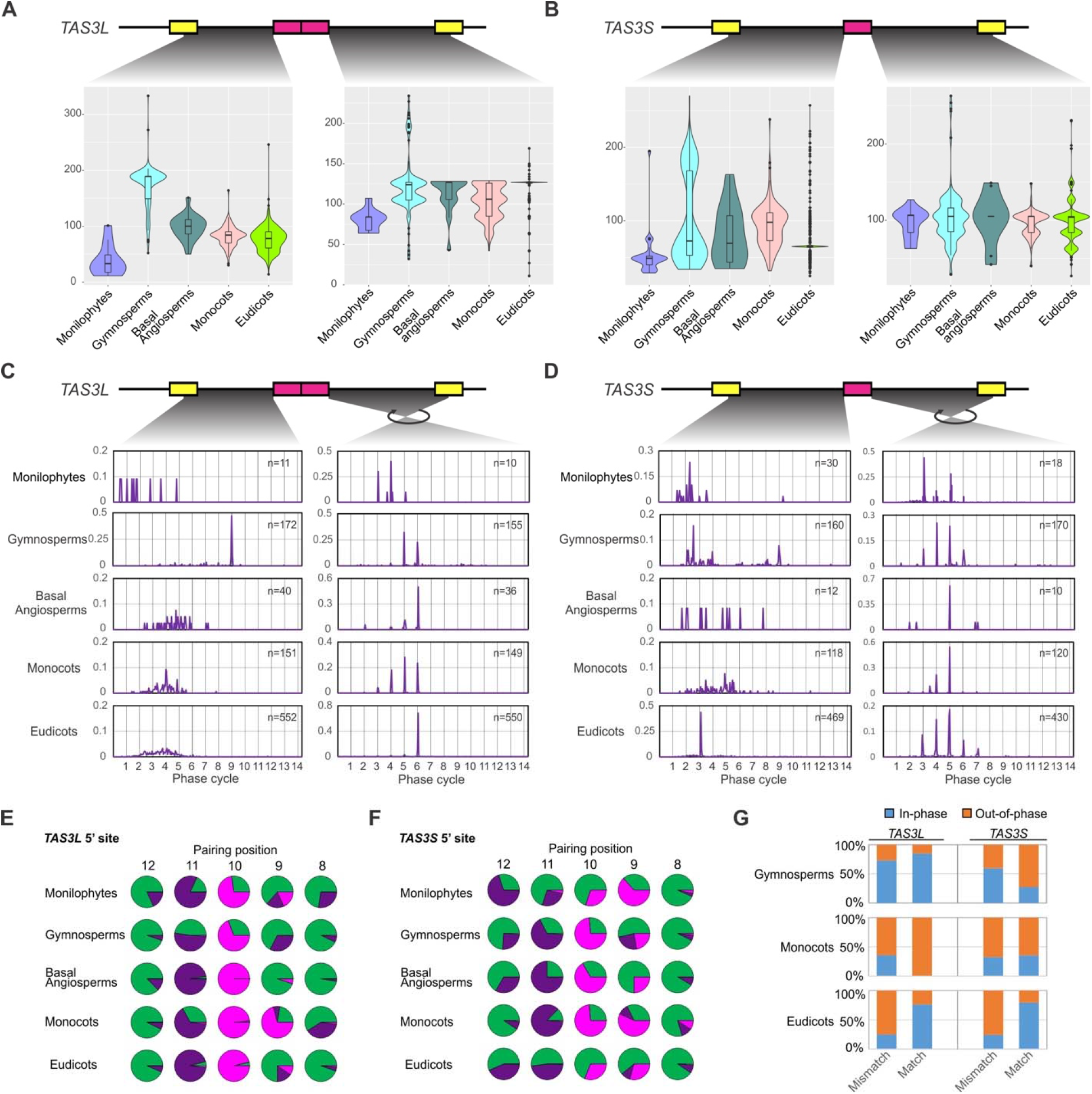
The distances between the two target sites of miR390 and the central tasiARF are under strong selection. Panels (A) and (B) display the variation of the distances between two miR390 target sites and tasiARF of *TAS3L* (panel A) or *TAS3S* (panel B) genes in different lineages of vascular plants. In both panels, the lower graphs contain violin plots for each lineage representing the distribution of these distances; internal boxes represent the median as a heavy line surrounded by a box defining the upper and lower quartiles. Panels (C) and (D) display the distribution of the distances between two miR390 target sites and tasiARF of *TAS3L* (C) or *TAS3S* (D) genes in different lineages of vascular plants. The Y-axis is the percentage of *TAS3* genes with distances occurring within a given position (the X-axis). The 21-nt phased positions (phase “cycles”) are marked as grey gridlines. Panels (E) and (F) display the variation in pairing of the 8^th^ to 12^th^ nucleotide positions (relative to the 5′ end of miR390) of the 5′ target site of *TAS3L* (E) and *TAS3S* (F). The type of miR390-*TAS3* pairing observed at different nucleotide positions, relative to the 5′ end of miR390, with A:U/C:G matches denoted in green, G:U matches in purple, and all mismatches in pink. (G) Ratio of the 5′ miR390 target sites in phase or out-of-phase to the tasiARF in terms of different nucleotide pairing at the 10^th^ position (match: U; mismatches: A, C, G as the 10^th^ nucleotide of miR390 is “A”).

Next, we assessed the distance from the tasiARFs to a miR390 cleavage site in terms of the phase cycles of phased siRNAs, to determine which site was the trigger. The tasiARFs of *TAS3L* are mostly out-of-phase with the 5′ target site, with the exception of those from gymnosperms in which the tasiARFs are consistently positioned at the 9^th^ cycles according to the cleavage site of the 5′ site (Fig. 5C left). In contrast, the *TAS3L* tasiARFs are consistently in phase to the 3′ site; in other words, the distances of the 3′-site → tasiARF were almost uniformly a multiple of 21 nucleotides, despite considerable length variation in some groups (Fig. 5C). For *TAS3S*, its tasiARF is largely not in phase to the 5′ site, except in the eudicots, which had a consistent 5′-site → tasiARF distance of approximately three cycles, or 65 nucleotides. As with TAS3L, although variation in the length was observed for the 3′site → tasiARF region, the distance was almost uniformly phased as well, i.e. a multiple of 21 nucleotides (Fig. 5D). These results indicated that the 3′ site is the main trigger site of tasiARF generation in both *TAS3L* and *TAS3S*, but the gymnosperm *TAS3L* and eudicot *TAS3S* likely also generate tasiARFs triggered by the 5′ miR390 target site.

### The cleavability of the 5′ site and its in-phase tasiARF are selected coordinately

The non-cleavable feature of 5′ miR390 target site is functionally important for its role as a binding site of the miR390-RISC complex, and this non-cleavability results from the presence of a mismatch at the 10^th^ position of the target site pairing (Montgomery et al. 2008b; Axtell et al. 2006). As aforementioned, our analysis of the middle region of the miR390:target-site pairing of the 5′ site (Fig. 4B) demonstrated that, consistent with previous studies, the 10^th^ position mismatch is indeed conserved in the majority of the *TAS3 genes* in vascular plants. However, we also observed that a not-insignificant fraction of interactions of the 10^th^ position of miR390 with *TAS3* are perfectly paired, especially in monilophytes, gymnosperms and eudicots (Fig. 4B). Given the finding that the tasiARF in gymnosperm *TAS3L* and eudicot *TAS3S* copies are mostly in phase with the 5′ site as well, it is conceivable that the portion of matched 10^th^ position is contributed by the 5′ sites capable of setting the phase of the tasiARF. To check this possibility, we separated the 5′ sites of *TAS3L* from those of *TAS3S*, and focused our analyses on the middle region (8^th^ to 12^th^ positions, relative to the miRNA), as shown in Fig. 5E and F. Although the general pattern was similar for *TAS3L* and *TAS3S*, i.e. a predominant 10^th^ position mismatch in most lineages and preferential 11^th^ position G:U pairing, we found a few dissimilarities between *TAS3L* and *TAS3S* in the pairing at these positions. Most noticeable was the level of perfect matches at the 10^th^ position for *TAS3S* compared to the majority of mismatches in *TAS3L* at the same position (Fig. 5E). We then asked whether those 5′ sites in phase to tasiARF were more likely to display a 10^th^ position perfect match or not. When we divided the 10^th^ position into two groups, the matched group (with a “U” matching the 10^th^ position “A” of miR390), and the mismatched group (“A”, “C”, or “G”), and we calculated the proportion of in-phase target sites and out-of-phase target sites, separately. The matched group had a much higher proportion of in-phase sites in eudicots (Fig. 5G), suggesting that the in-phase and cleavable 5′ site was coordinately selected during *TAS3* evolution in eudicots.

### Strong mutual selection between tasiARF and its target site in ARF genes

The miR390-*TAS3-ARF* pathway exerts its function via the silencing of a subgroup of *ARF* genes, *ARF2/3/4* in Arabidopsis (Allen et al. 2005). In Arabidopsis, the *ARF* genes are classified into three clades Clade A (*ARF5/6/7/8*); Clade B (*ARF1/2/3/4/9*), and Clade C (*ARF10/16/17*) (Finet et al., 2012). The vascular tasiARFs target *ARF2/3/4* belong to Clade B, the ancestor of which likely emerged in liverworts (Finet et al. 2013). Typically, *ARF2* has a single target site for the tasiARF, and *ARF3/4* have two target sites (Allen et al. 2005; Axtell et al. 2006). As described above, the *ARF* genes in Clade B of bryophytes are regulated by tasiARF-a1 to -a3, thereafter in evolution, this group was targeted by the tasiARF that emerged in vascular plants. However, we found that some Clade B genes from mosses (for example, from *P. patens*) bear analogous target site sequence of the vascular tasiARF, suggesting that this target site predates the emergence of the tasiARF of vascular *TAS3* genes (Fig. S6). Combining these data with the *ARF* evolution history illustrated in Finet et al. (2012), we summarized the likely path of diversification of tasiARF target sites during the evolution of the Clade B *ARF* genes (Fig. 6A). The interaction pattern of tasiARF with *ARF* genes was likely formed in lycophytes with only one target site. In Monilophytes, genes in Clade B are targeted at a single site in most species, but a few species display dual target sites. Thereafter, in evolutionary terms, this dual targeting was maintained in the subclade, and likely eventually gave rise to the *ARF3/4* genes, while the single targeting was selectively retained in the *ARF2* subclade, but lost in the *ARF1/9* group (Fig. 6A).

**Figure 6.**
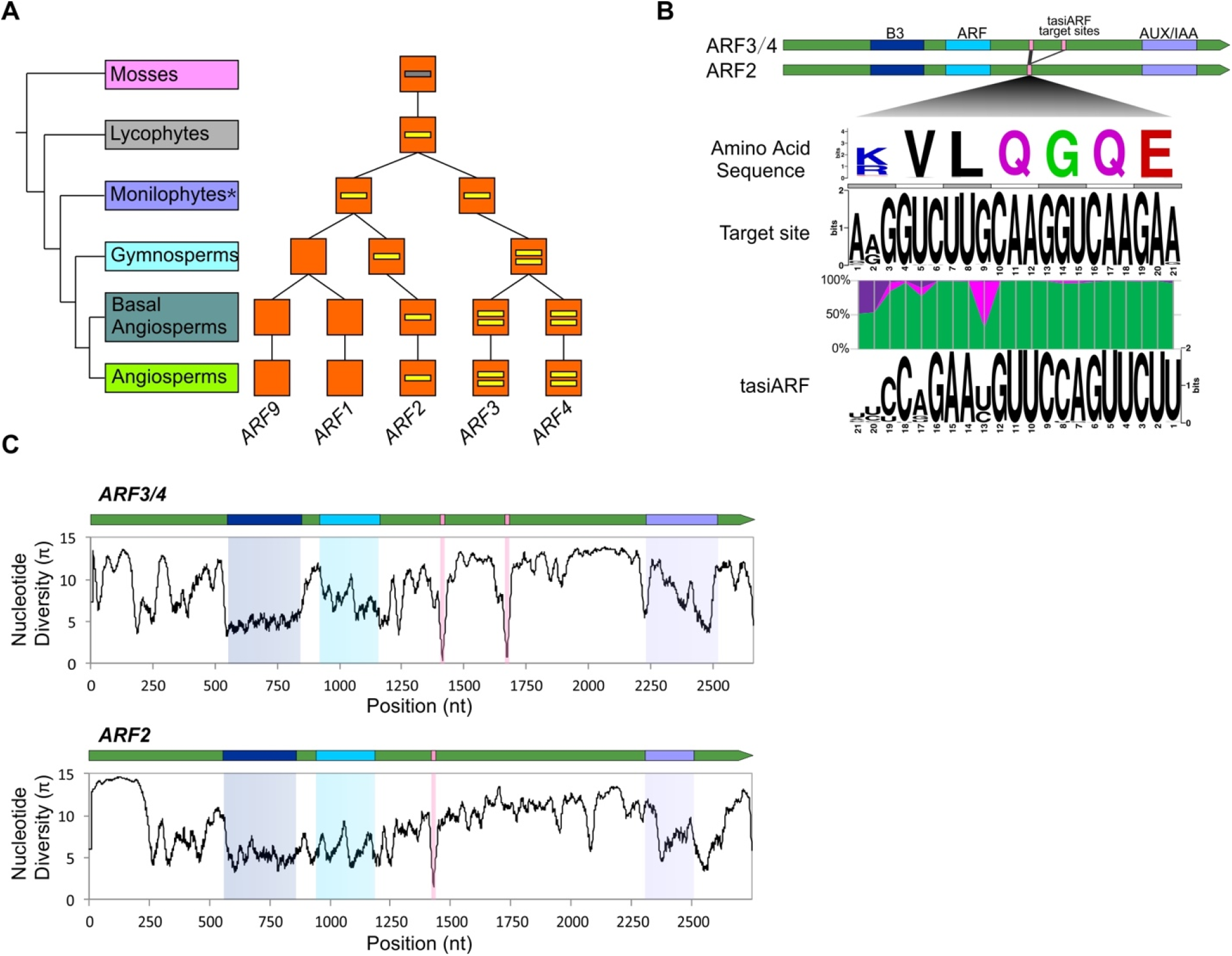
Evolutionary diversification of *tasiARF* target sites in *ARF* genes. (A) Evolution of the number of *tasiARF* target sites in plant *ARF* genes. The evolutionary route of *ARF* genes was adapted from Finet *et al.* (2012). The number of short yellow lines in orange boxes denote the number of tasiARF target sites. The grey line means that there are potential tasiARF target sites in *ARF* genes in mosses. In monilophytes (marked with a “*”), some *ARF3/4* homologous genes have already evolved two tasiARF target sites. (B) Sequence features of the target site of tasiARF in *ARF* genes and their encoded proteins. Gene structures of tasiARF-targeted *ARF2/3/4* are displayed on the top, including the encoded protein motifs, with the tasiARF target site indicated as pink bars. The target site encodes a short peptide with a consensus sequence of K/RVLQGQE, as indicated with the encoding sequence. Pairing between tasiARF and its target site is color-coded with A:U/C:G matches denoted in green, G:U matches in purple, and all mismatches in pink. (C) Distribution of nucleotide diversity along tasiARF-targeted *ARF2/3/4* genes, with the encoded functional domains and tasiARF target site marked in colors according to those in panel B.

The target sites of vascular tasiARF were located in the middle region between two functional domains (ARF and AUX/IAA) of *ARF2/3/4* genes (Fig. 6B). We recently reported that the miR482/2118 family displays significant sequence variation at positions matching the 3^rd^ nucleotide of codons at the miRNA target site, implying a strong selection from the functionally important P-loop motif of NB-LRR proteins that shapes miRNA-target pairing (Zhang et al. 2016). In contrast, the tasiARF sequence is of much lower sequence divergence, and it did not show a pattern like the miR482/2118 family, indicating the selection on tasiARF pairing is distinct from the miR482/2118 case. Similarly, the tasiARF target sites, unlike the miR390 target sites in *TAS3* genes which are of considerable diversity, are less divergent in sequence, and consistently encode the amino acid sequence K/RVLQGQE (Fig. 6B). We also assessed nucleotide diversity of the *ARF2/3/4* genes and we found that the three functional domains were, as expected, of relatively low nucleotide diversity. However, the tasiARF target sites (one in *ARF2* and two in *ARF3/4*) showed substantially lower nucleotide diversity, even compared to the encoded, conserved functional domain, indicating a strong selection on them during evolution (Fig. 6C). Given the fact that tasiARF sequences in *TAS3* genes are also extremely conserved in vascular plants, we hypothesize that there is strong mutual selection between tasiARF in *TAS3* genes and its target sites in *ARF* genes.

### New regulatory mechanism of TAS3 genes

When we were annotating *TAS3* genes, we observed several other previously-undescribed features of the *TAS3* gene family. First, we found a few vascular *TAS3* genes that display transcript isoforms generated by alternative splicing. A good example is the *TAS3c* locus in maize, in which many alternative splicing sites giving rise to numerous transcript isoforms (Fig. 7A). These isoforms selectively spliced out the three essential components of the *TAS3* gene; for instance, the 5′ site is missing for splice variant T4, the 3′ site is missing for T1 and T2, both target sites missing for T5, T8 and T9, and all of the target sites and tasiARFs missing for T7, T10 and T13 (Fig. 7A). While we currently have no evidence of functional relevance for these variants, it’s possible that alternative splicing can serve as another layer of regulation to fine-tune the activity of *TAS3* genes and subsequent tasiARF production. For example, there is evidence that small ORFs encoded by *TAS* genes play functional roles (Yoshikawa et al., 2016), and these splice variants could mediate ribosome loading, stalling, or peptide production, independent of tasiRNA biogenesis.

**Figure 7.**
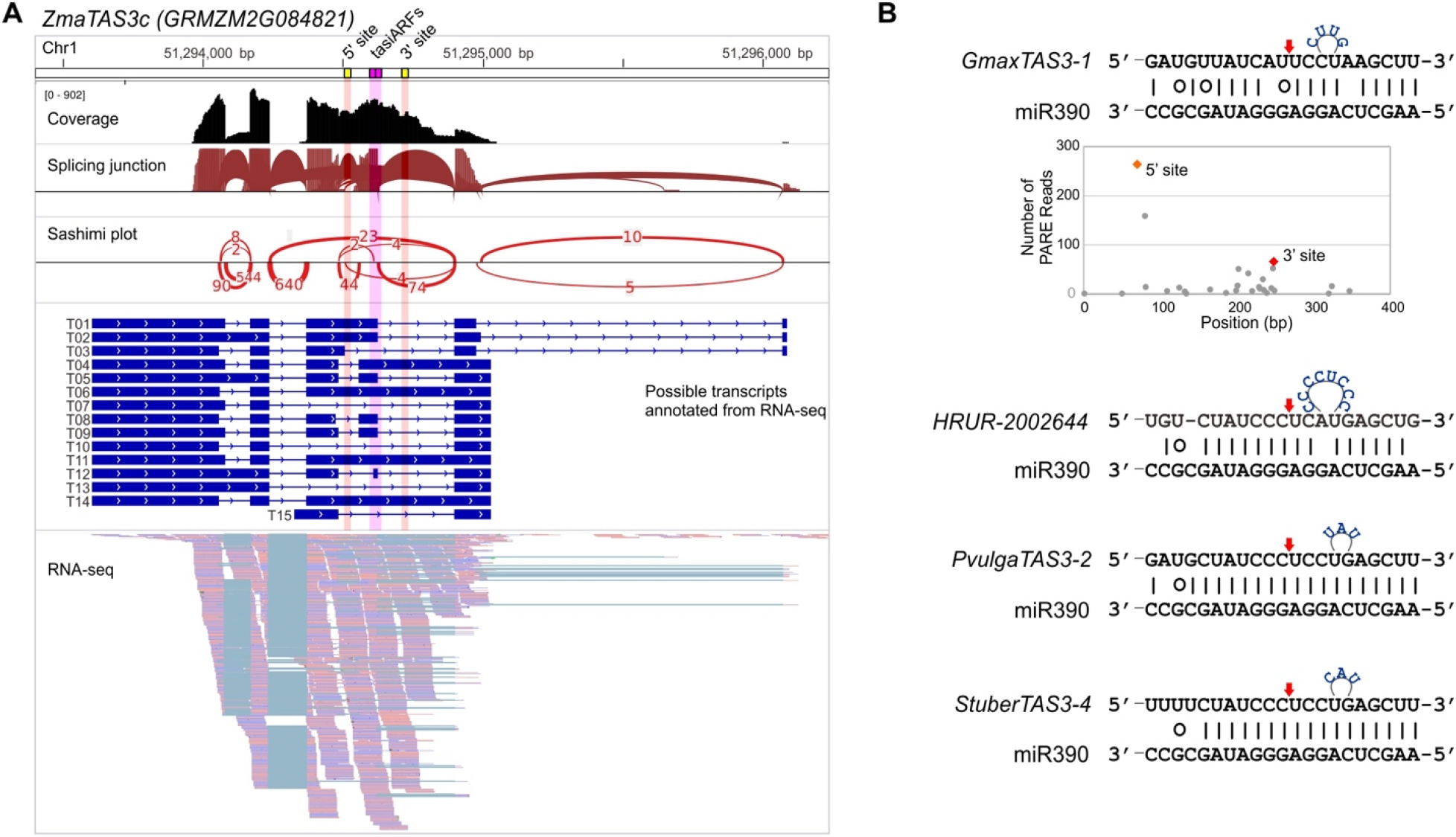
New regulatory features found for the miR390-*TAS3-ARF* pathway. (A) Alternative splicing affects the structure of *TAS3* transcripts in maize. tasiARF (pink) and the two target sites of miR390 (yellow) are marked at the top, and they are alternatively present/absent in different splicing variants (T1…T15). The “Sashimi plot” displays the frequency with which different splice ends were joined in the RNA-seq data (at bottom). (B) miR390-interactions predict a large bulge in the so-called “seed” region of the miRNA:target interaction, for the 3′ target site of several *TAS3* genes; the example at the top is from soybean. Soybean PARE data were consistent with the successful cleavage of this 3′ target site of miR390 in *GmaxTAS3-1*. Three other cases from different species are displayed below this, from *Utricularia sp.* (*HRUR-2002644*), common bean (*Phaseolus vulgaris, PvulgaTAS3-2*), and potato (*Solanum tuberosum, StuberTAS3-4*).

Another new feature that we observed is an abnormal pairing interaction of miR390 with a few target sites in *TAS3* genes; this pairing displays large bulges in the seed sequence region (2^nd^ to 13^th^ positions) (Fig. 7B). The first example is the *TAS3-1* gene in soybean (*Glycine max*). We can infer based on the register of the siRNAs that its 3′ target site is cleaved to set the phasing, despite a predicted 4-nt bulge present between the 6^th^ and 7^th^ positions of miR390 (Fig. 7B). Another three cases were predicted to generate an 8- or 3-nt bulge (Fig. 7B). This type of abnormal, bulge-containing miRNA-target interaction was recently reported and validated in Arabidopsis for miR398 (Brousse et al. 2014), the only other known case of this type. Our results suggest that this is pairing is not unique to miR398, and large asymmetrical bulges in miRNA:target pairing are at least allowable in plants.

## Discussion

The presence of miR390 and *TAS3* was tracked back to liverworts (Lin et al. 2016), while the ARF domain encoded by *ARF* genes likely first appeared in the land plants (Finet et al. 2013). We demonstrated that *TAS3* in liverworts produces tasiRNAs to target *ARF* genes, suggesting this was the earliest function of *TAS3*, a key function maintained throughout land plants. We also observed in liverworts the conservation of another *TAS3*-derived tasiRNA that, in mosses, targets *AP2* genes (referred to as tasiAP2), but we were unable to confirm this function in liverworts. It is possible that this tasiRNA in liverwort *TAS3* genes emerged before the appearance of tasiAP2 target sites in *AP2* genes, or this tasiRNA has an unidentifiable function or target.

Although the role of *TAS3* in regulating *ARF* genes is conserved across land plants, the bryophyte *TAS3* genes are structurally different from those in vascular plants (Axtell et al. 2006). In other words, tasiARFs are different in sequence between bryophytes and vascular plants. In our model for tasiARF evolution, the tasiARF was derived from the duplication of the 5′ target site of miR390, and the short *TAS3* variant (*TAS3S*) is the ancestor of the long *TAS3* (*TAS3L*). We identified the vascular *TAS3* in a lycophyte, indicating the transition of tasiRNA sequences is likely associated with the development of vascular tissue in plants, as lycophytes were among the first vascular plants on earth. Measured across the vascular plants, there are nearly always two types of *TAS3* genes (*TAS3S* and *TAS3L*) present in each plant genome and totaling approximately four members in most species. The deep conservation of these structures suggests they are not functionally redundant. Future work could address why, perhaps by selective deletion of the two types using CRISPR/Cas9. Another striking observation was that the *TAS3* copy number is significantly expanded in conifers, reminiscent of the expansion of *NB-LRR-*targeting miRNAs (Xia et al., 2015). Despite evidence of whole genome duplications in spruce (Li et al. 2015), the >10-fold higher copy number in conifers relative to angiosperms is extraordinary. Perhaps functional differences in tasiRNA movement in gymnosperms required added copies of *TAS3* genes, but these copies were made redundant and lost by angiosperm-specific evolutionary adaptations, possibly improved tasiRNA mobility.

In plants, miRNA/miRNA duplexes are released by two sequential cuts of their hairpin precursors by DCL1; these cuts can occur either base-to-loop or loop-to-base (Bologna et al. 2009, 2013). For other miRNAs, it was observed that a conserved length of the basal stem region ensures accurate cuts made by DCL1 (Werner et al. 2010; Song et al. 2010; Mateos et al. 2010). We found that miR390 is processed in a base-to-loop direction, with the first cut by DCL1 occurring at a position ~15 nt from a basal unpaired region (> 4 nt). While this basal region (the unpaired region to the site of the first cut) is a length consistent with other base-to-loop processed miRNAs, by comparison across the seed plants, we found that the sequence is also relatively conserved, indicating that selection can maintain bases in the hairpin other than the miRNA/miRNA*. The conservation of this paired region was recently described for many plant miRNAs (Chorostecki et al. *in preparation*).

One of the major differences between two main mechanisms of tasiRNA/phasiRNA biogenesis (“one-hit” and “two-hit” models) is the direction of tasiRNA production. In the “two-hit” model, tasiRNAs are produced in a 3′ to 5′ direction, in contrast to the predominant 5′ to 3′ Dicer processing (i.e. the “one-hit” model). miR390-*TAS3* is the quintessential “two-hit” locus, yet its 3′ to 5′ processing is distinctive and rare. Evolutionary analyses of miR390-*TAS3* pairing revealed two distinct patterns of pairing of the two target sites: (i) a conserved mismatched region mainly caused by the 10^th^ position of the 5′ site (previously known – see below), and (ii) an open, unpaired region in the 3′ end of the 3′ target site (from our study). The wide conservation in vascular plants of these features implies functional relevance. Studies in Arabidopsis have shown that the non-cleavability of the 5′ site, caused by the central mismatch (10^th^ position), is essential, mediating miR390 binding via AGO7 (Rajeswaran and Pooggin 2012). Changing the 10^th^ mismatch into a perfect match compromises tasiRNA biogenesis (Axtell et al. 2006; Montgomery et al. 2008a). However, a substantial portion of *TAS3* genes, especially the *TAS3S* subset, having a cleavable 5′ site (A:U pair at the 10^th^ position), and many of these sites, particularly in eudicots, trigger tasiARF production. This indicates that the non-cleavability of the 5′ site is helpful but not necessary for tasiARF production. Another notable feature of the 5′ site pairing is the predominant G:U pairing at the 11^th^ position; this preferential wobble pairing might be helpful for maintaining the non-cleavability of the 5′ site, which is believed to be mainly caused by the 10^th^ position mismatch. In contrast, the pairing of the 3′ site has a consistently matched middle region, but an open, unpaired 3′ end region. The paired middle region could ensure the cleavage of the 3′ site, make it the typical trigger site for secondary tasiRNAs. The 3′ end open region may direct the 3′ to 5′ production of *TAS3* tasiRNAs. Perhaps after cleavage, the 3′ end open region makes the cleaved mRNA end more accessible to RDR6 to facilitate downstream tasiRNA production.

Besides the cleavability of the target site, the distance between the miR390 target site and tasiARF also appears to be a determinant for phasiRNA biogenesis. tasiARF production requires distances in multiples of 21 nucleotides from the cleavage site (“in register”). We showed that the distance of the 3′ side of *TAS3* is more consistently a multiple of 21 nt despite considerable length variation; the 3′ site also displayed fewer 10^th^ position mismatches (i.e. better cleavability). However, we noted several exceptions. The *TAS3L* in gymnosperms and the *TAS3S* in eudicots had a highly consistent distance on the 5′ side, in approximate phase with tasiARF, and the cleavability of the 5′ site is coordinately selected with the in-phase distance to the tasiARF in eudicots, suggesting that the 5′ site in those *TAS3* genes is likely to serve as a trigger site of tasiARF production as well. Therefore, our results suggest that some *TAS3* loci in vascular plants are likely bi-directionally processed, consistent with the observation of the original bidirectional processing of functional tasiRNAs in bryophytes (Axtell et al. 2006). For instance, the two target sites of *TAS3* in *P. patens* are both cleavable, and tasiARF-a2 is in phase with the 3′ site while tasiARF-a3 is in phase with the 5′ site. This bidirectional processing thus yields additional questions about this “two-hit” mechanism. How is the activity of the two sites coordinated? Does cleavage occur simultaneously at both sites or one site at a time?

miRNAs, tasiRNAs, or other type of sRNAs and their targets genes are a pair of partners, functioning via their interactions, based on sequence complementarity. Few studies have deeply investigated this sRNA:target partnership over evolutionary time. In the case of the widely conserved miR482/2118 family, we described selection from target protein-coding genes to miRNAs; in other words, the essential function of the P-loop encoded in *NB-LRR* genes, targeted by miR482/2118, is most important, as miRNA variation matches a degenerate nucleotide change at the third position of each codon in the target (Zhang et al. 2016). In the current study, we detected a distinct pattern of selection between tasiARF and target site in *ARF* genes in vascular plants. Both are depleted of variation, indicating a strong mutual selection. The tasiARF target site sequences in *ARF* genes show no periodical variation (at the third position), indicating the target site sequence is not under strong selection at the amino acid level, in accordance with the location of the tasiARF target site between two encoded domains of ARF proteins, the ARF domain and AUX/IAA domain. The target site in the middle region is of less functional importance at the protein level. This is in contrast to the location of miR482/2118 target site in a functionally critical domain (Zhang et al. 2016). However, the sequence variation (nucleotide diversity) of the tasiARF target sites in *ARF* genes is dramatically less than other gene regions, even the conserved functional protein domains, suggesting that the tasiARF target site is under a selective force stronger than selection than that of the encoded protein domains. Combined with the fact that tasiARF sequences in vascular *TAS3* copies demonstrate substantially less sequence variation, we believe that there is a robust selective connection between tasiARF and its target site in *ARF* genes, which permit little sequence variation in either component over evolutionary time.

## Methods

### Genome sequences and 1KP data

Genome sequences of 159 species were retrieved from either the Phytozome or NCBI. The assembled transcriptome data of the 1000 Plant Transcriptome Project (“1KP”) was kindly shared by the Wang lab at the University of Alberta, Canada (Matasci et al. 2014).

### NGS data and analyses

RNA of *Phylloglossum drummondii* was extracted using PureLink Plant RNA Reagent. A sRNA library was constructed using the Illumina TruSseq sRNA kit, and sequenced on the Illumina HiSeq platform at the University of Delaware. The sRNA data was deposited in NCBI GEO (Gene Expression Omnibus) under the accession number GSE90706.

sRNA and PARE data of *Marchantia polymorpha* were retrieved from NCBI Short Read Archive (SRA) under accession numbers SRR2179617 and SRR2179371, respectively (Lin et al. 2016). sRNA reads were mapped to reference genome or transcripts by Bowtie (Langmead et al. 2009), and PARE data was analyzed using Cleaveland 2.0 (Addo-Quaye et al. 2009).

Paired-end RNA-seq data for maize were downloaded from NCBI SRA under accession numbers SRR1213570 and SRR1213571 (Wang et al. 2015). RNA-seq reads were mapped to maize genome using STAR v2.4.2a (Dobin et al. 2013) and transcripts of the *ZmaTAS3c* locus were annotated using Cufflinks v2.2.1 (Trapnell et al. 2012). The mapped bam files of two libraries were merged and viewed using Integrative Genomics Viewer v2.3.59 (Robinson et al. 2011).

### Homologous gene identification

For the identification of *MIR390* genes, mature sequences of miR390 were retrieved from miRBase, and used to search for homologous sequence using FASTA36 allowing two mismatches. After that, ± 500 bp sequence were excerpted for each homologous sequence from reference sequences and used for the evaluation of secondary structure. Only those genomic loci or transcripts with a good stem loop structure (≤ 4 nt mismatches and ≤ 1 nt bulge) and with the mature miRNA in the 5′ arm were regarded as good *MIR390* genes.

For the identification of *TAS3* genes, < 500 bp genomic loci (for genomes) or EST sequences (for transcriptome data) with evidence of at least two components of the two miR390 target sites and one tasiRNA (tasiARF for vascular plants, tasiAP2 or tasiARF-a2 for bryophytes) were considered as *TAS3* candidates. Their identity as a *TAS3* gene was further assessed by manual sequence comparisons. The tool MEME (Bailey et al. 2009) was also used to profile the signature sequence motif of *TAS3* genes.

To identify tasiARF-targeted *ARF* genes, firstly Arabidopsis and subsequently rice ARF proteins were used as bait sequences to identify *ARF* homologous genes, using either TBLASTN for annotated genomes or 1KP transcriptome data or genBlast (She et al. 2011) for unannotated genomes. Secondly, TargetFinder (https://github.com/carringtonlab/TargetFinder) was used to identify tasiARF-targeted *ARF* genes. Thirdly, *ARF3/4* and *ARF2* genes were distinguished by the number of target sites as *ARF3/4* genes have two tasiARF target sites and *ARF2* genes have only one target site. AGO proteins were identified using BLASTP for selected annotated genomes or TBLASTN for 1KP data using Arabidopsis and rice AGO proteins as bait sequences. Only full-length AGO protein sequences from sequenced genomes and AGO sequences with ≥ 800 amino acids from the 1KP data were chosen for subsequent phylogenetic tree construction.

### Multiple alignment and tree construction

Amino acid sequences of Argonautes (≥ 800 amino acids), annotated from transcripts and genomes, were aligned using MUSCLE v3.8.31 with default parameters (Edgar 2004). The regions poorly aligned were trimmed using trimAl v1.4 (Capella-Gutiérrez et al. 2009), and the trimmed alignments were used for construction of a maximum likelihood (ML) tree using RAxML v8.1.1 under the GTRCAT model (Stamatakis 2014). For a tree of bryophyte *TAS3* genes (Fig. S3B), the nucleotide sequences of those genes were aligned and edited similarly, and the ML tree was made using RAxML under the PROTGAMMAAUTO model. For each tree, 100 replicates were conducted to generate bootstrap values. The trees were viewed using Dendroscope v3.5.7 (Huson and Scornavacca 2012).

Jalview was used for the viewing of alignment results (Waterhouse et al. 2009). The R package was used to make violin plots and conduct statistical analyses. Sequence logos of sRNA and target sites were generated using Weblogo (Crooks et al. 2004). To calculate the nucleotide diversity (π) of *ARF* genes, the amino acid sequences of *ARFs* were generated by translation of the genes, aligned using MUSCLE, then the protein sequence alignment was used to generate the alignment of nucleotide sequences using PAL2NAL (Suyama et al. 2006). Subsequently, poorly aligned regions, those with <30% nucleotide coverage, were removed, and finally the nucleotide diversity (π) at a single nucleotide level was calculated using a 20 nt sliding window.

## Acknowledgements

We thank members of the Meyers lab for helpful discussions and input. This study was supported by US National Science Foundation, Division of Integrative Organismal Systems award #1257869 and the Chinese Thousand Young Talents Program. We are grateful to Dennis Stevenson and Ryan Lister for assistance in obtaining *Phylloglossum drummondii* material. We are also grateful to Gane Ka-Shu Wang for assistance with access to the 1KP data.

